# Trait Reward Sensitivity Modulates Connectivity with the Temporoparietal Junction and Anterior Insula during Strategic Decision Making

**DOI:** 10.1101/2023.10.19.563125

**Authors:** Daniel Sazhin, James B. Wyngaarden, Jeff B. Dennison, Ori Zaff, Dominic Fareri, Michael S. McCloskey, Lauren B. Alloy, Johanna M. Jarcho, David V. Smith

## Abstract

Many decisions happen in social contexts such as negotiations, yet little is understood about how people balance fairness versus selfishness. Past investigations found that activation in brain areas involved in executive function and reward processing was associated with people offering less with no threat of rejection from their partner, compared to offering more when there was a threat of rejection. However, it remains unclear how trait reward sensitivity may modulate activation and connectivity patterns in these situations. To address this gap, we used task-based fMRI to examine the relation between reward sensitivity and the neural correlates of bargaining choices. Participants (N = 54) completed the Sensitivity to Punishment (SP)/Sensitivity to Reward (SR) Questionnaire and the Behavioral Inhibition System/Behavioral Activation System scales. Participants performed the Ultimatum and Dictator Games as proposers and exhibited strategic decisions by being fair when there was a threat of rejection, but being selfish when there was not a threat of rejection. We found that strategic decisions evoked activation in the Inferior Frontal Gyrus (IFG) and the Anterior Insula (AI). Next, we found elevated IFG connectivity with the Temporoparietal junction (TPJ) during strategic decisions. Finally, we explored whether trait reward sensitivity modulated brain responses while making strategic decisions. We found that people who scored lower in reward sensitivity made less strategic choices when they exhibited higher AI-Angular Gyrus connectivity. Taken together, our results demonstrate how trait reward sensitivity modulates neural responses to strategic decisions, potentially underscoring the importance of this factor within social and decision neuroscience.

## Introduction

Social situations such as negotiations often require people to strategically consider social norms while minimizing the threat of being rejected. It is understood that people act fairly when they could be rejected in the Ultimatum Game (UG; Güth et al., 1982; Wells & Rand, 2013) and selfishly when there is not a threat of rejection in the Dictator Game (DG; Engel, 2011; Kahneman et al., 1986). Thus, people exhibit strategic behavior by making smaller contributions in the DG than in the UG (Charness & Gneezy, 2008). Past investigations suggested there are relations between strategic behavior and measures of social functioning such as emotional intelligence (Kench et al., 2007) and Machiavellianism (Spitzer et al., 2007). A possible explanation for strategic behavior is the social heuristics hypothesis, which suggests people share more or less intuitively based on self-interest, and greater deliberation yields more strategic choices (Rand, 2016; Rand et al., 2016).

Strategic decisions as defined by making larger contributions in UG compared to DG have also been associated with brain activation in the ventral striatum (VS), dorsal lateral prefrontal cortex (dlPFC), and lateral orbitofrontal cortex (OFC) (Spitzer et al., 2007). Other work has implicated dorsal anterior cingulate cortex (dACC) and the posterior cingulate cortex (PCC) in strategic decision making (Weiland et al., 2012). Decisions made in social contexts reliably elicit activation in the right temporoparietal junction (rTPJ) (Behrens et al., 2008; Carter et al., 2012; Dennison et al., 2022), and higher rTPJ activation is associated with greater contributions in the DG (Gianotti et al., 2018; Morishima et al., 2012). Further, stimulation of the right dlPFC is associated with proposing greater contributions in UG and less in DG (Knoch et al., 2006; Ruff et al., 2013; Strang et al., 2015). Finally, people make lower contributions in the DG after stimulation the right dlPFC (Zinchenko et al., 2021). In sum, there is evidence that brain activation can distinguish between some strategic decision making in social contexts.

Relatively less is known, however, about how strategic decisions in bargaining situations are modulated by task-dependent changes in connectivity across neural circuits supporting reward related decision-making and social cognition (Friston et al., 1997). Past research suggests that signals related to the receipt of rewards are encoded through corticostriatal connectivity (D. V. Smith, Rigney, et al., 2016). Moreover, VS-TPJ connectivity (Park et al., 2017) and dorsal striatum-lateral PFC connectivity (Crockett et al., 2017) were modulated by contributions proposed in DG. Since past findings suggest that anticipating the intentions of another person in an investment game (Zhu et al., 2012) and greater contributions in UG versus DG (Spitzer et al., 2007) were associated with elevated VS responses, it is possible that corticostriatal connectivity may be modulated by social contexts in bargaining situations.

Additionally, individual differences in trait reward sensitivity may affect how people make social valuations, possibly moderating neural connectivity in social contexts. Reward sensitivity has been studied in clinical contexts (Alloy et al., 2016; Carver & White, 1994; Nusslock & Alloy, 2017), revealing that people who are hyper and hyposensitive to rewards are at risk for substance use and bipolar or depressive disorders (Bart et al., 2021). However, little is known about how corticostriatal connectivity is modulated by reward sensitivity (Sazhin et al., 2020). For instance, people who are more sensitive to rewards may overvalue their initial endowment in UG and DG contexts and may be loath to share it with a stranger.

Since reward sensitivity is associated with risky behavior (Scott-Parker & Weston, 2017), higher Machiavellianism (Birkás et al., 2015), and with more strategic behavior (Scheres & Sanfey, 2006), it is plausible that mechanisms underlying strategic decision making may be modulated by reward sensitivity through VS activation or elevated task-based connectivity with the VS. Evidence supporting this interpretation would suggest that strategic decisions may be mechanistically driven by reward processing and that reward sensitivity is a reflection of bottom-up reward responses. Alternatively, strategic decisions may evoke cognitive processes involved in attention and social decision making from brain regions such as the TPJ. Evidence supporting this interpretation would suggest that strategic decisions are driven by top-down cognitive processes and may be modulated by trait reward sensitivity. Overall, examining the role of reward sensitivity and brain responses during strategic decisions could unpack reward, attentional, or value-based decision-making mechanisms that facilitate overcoming social heuristics to act on self-interest.

Since the VS is sensitive to social valuation (Chein et al., 2011; Fareri & Delgado, 2014), it is plausible that trait reward sensitivity may modulate VS response to social contexts. Testing these relations could help unravel how aberrant reward processing promotes maladaptive decisions that contribute to substance use (Dalley & Robbins, 2017), or possibly diminishes strategic behavior in social situations. Thus, our aims in this investigation were to assess how brain activity and connectivity are modulated by one’s strategic decisions, and the extent to which these relations vary by trait reward sensitivity. Using functional magnetic resonance imaging (fMRI), we administered Ultimatum and Dictator Games to participants to investigate associations between strategic behavior, reward sensitivity, and brain connectivity. The study examined activation patterns during both endowment and decision phases, corticostriatal connectivity during the decision phase, and how these patterns were modulated by strategic behavior and reward sensitivity.

To examine these questions, we assessed several pre-registered hypotheses (https://aspredicted.org/55gd8.pdf). Participants proposed offers in DG and UG (eg: DG-P and UG-P) and received offers as a recipient (UG-R). We expected greater activation of the VS and vmPFC during the endowment of money, and that reward sensitivity would potentiate activation in the VS and vmPFC. Such activation during endowment would suggest that reward receipt is modulated by reward sensitivity. Next, we investigated activation within the dlPFC, ACC, SPL, IPS, vmPFC, VS, and TPJ during each task condition and specifically in response to strategic decisions (UG-P > DG-P). We hypothesized that the dlPFC would exhibit stronger activation in response to strategic decisions. These findings during the decision phase would suggest that changes in social context, with respect to norm compliance, evoke differential activation in the brain. Finally, we expected to find elevated ventral striatal responses to strategic behavior (UG-P > DG-P) during the decision phase to be associated with enhanced task-dependent changes in connectivity in regions modulated by social information (e.g., vmPFC, mPFC, and TPJ). In addition, we hypothesized that these neural effects would be enhanced in individuals with higher level of self-reported reward sensitivity. Such findings would suggest that reward sensitivity is an important dimension of understanding brain responses associated with strategic behavior.

Our analyses focus on two key questions. First, how do strategic decisions in social situations modulate brain activation and connectivity? Second, how does trait reward sensitivity modulate brain connectivity while making strategic decisions? Assessing neural connectivity during strategic decision making and how reward sensitivity modulates these processes would 1) improve our understanding of the mechanisms of how people cooperate and defect in social situations, and 2) help determine how aberrant patterns of reward sensitivity may be a risk factor for maladaptive social decision making.

## Materials and Methods

### Participants

Although in our pre-registration (https://aspredicted.org/55gd8.pdf) we specified that imaging data would be collected from 100 participants (ages 18-22) (Sazhin et al., 2020), we ultimately recruited 59 participants (D. V. Smith et al., 2024) due to constraints imposed by the COVID-19 pandemic. Five participants were excluded from our neuroimaging analyses based on our pre-registered criteria and missing data. Specifically, three participants were excluded due to failure to respond during behavioral tasks, where there were greater than 20% missing responses on a given run. One participant was excluded due to incomplete behavioral data. One participant was excluded due to issues with data collection. Three of the 54 participants had one of the two task runs excluded due to excessive head motion. Our final neuroimaging sample resulted in 54 participants (mean age: 20.95 years, SD: 1.78 years; 24.1% male). Our final sample size (N = 54) would enable us to detect medium effects strategic behavior or reward sensitivity (f^2 = 0.15) or medium to large interaction effects (f^2 = 0.19) with 80% power and an alpha of 5%.

Several behavioral analyses related to social functioning had a more limited sample due to missing data. Specifically, 9 participants were missing behavioral data related to social functioning, resulting in a sample of 45 participants (mean age: 20.74 years, SD: 1.54 years; 24.4% male) for several behavioral analyses. All participants were compensated at a rate of $25 per hour inside the scanner and $15 per hour outside the scanner, and received bonuses based on their decisions, resulting in a total payment ranging from $140 to $155. Participants were recruited using Facebook advertisements and fliers posted around the Temple University campuses. We verified that participants were eligible to be scanned using fMRI by the following criteria: a) not being pregnant, b) free of major psychiatric or neurologic illness, and c) not under the influence of substances as evidenced by a breathalyzer test and urine drug screen. All the participants provided written informed consent as approved by the Institutional Review Board of Temple University (protocol number: 24452). Data was acquired using a 3T Siemens PRISMA MRI scanner at Temple University using the Ultimatum and Dictator Games.

### Procedure

Potential participants were identified based on their responses to an online screener questionnaire using the SONA research platform that assessed reward sensitivity using the Behavioral Activation Subscale (BAS; Carver & White, 1994) and the Sensitivity to Reward subscale (SR; Torrubia et al., 2001). Using methods consistent with our prior work (e.g., Alloy, Bender, et al., 2009), we compared results between both SR and BAS to ensure that participants were responding consistently and truthfully by excluding participants with scores that were less than +/-1 quintile on both subscales. Participants also were called on the phone and asked to abstain from alcohol or drug usage for 24 hours prior to the scan. Participants were excluded if they reported that they took any psychoactive medications. Participants attended two appointments, consisting of a battery of psychometric surveys, and a mock scan, followed by a second appointment consisting of the fMRI scan and behavioral tasks.

### Individual Difference Measures

#### Reward Sensitivity

To measure reward sensitivity, we used the Behavioral Activation Scale (BAS; Carver & White, 1994) and the Sensitivity to Punishment/Sensitivity to Reward Questionnaire Reward subscale (SPSRWD; Torrubia et al., 2001)). The BAS is a 20-item self-report questionnaire that measures sensitivity to appetitive motives. The SPSRWD is a 24-item self-report measure that assesses how people feel in response to rewarding stimuli.

#### Substance Use

Given the relation between reward sensitivity and substance use (Bart et al., 2021), it was important to control for alcohol and drug use disorders in all analyses that include reward sensitivity. To measure substance use, we used the Alcohol Use Disorders Identification Test (AUDIT; Babor et al., 1992) and the Drug Use Identification Test (DUDIT; A. Berman et al., 2003; A. H. Berman et al., 2005). The AUDIT is a 10-item self-report measure that assesses frequency of usage over the past year and the self-reported extent to which alcohol use affects the person’s life. The DUDIT scale is an 11-item self-report measure counterpart of the AUDIT that assesses frequency and disruptiveness of non-alcohol related substance use. DUDIT contains references to a wide array of substances, including marijuana, cocaine, and others.

#### Social Functioning

To measure social functioning, we measured trait emotional intelligence and attitudes toward rejection. The trait Emotional Intelligence (EI) questionnaire (TEIQe) is a 30-item self-report measure that assesses individual differences in trait empathy, emotion regulation and perspective taking in emotional contexts (Petrides, 2009). Attitudes toward reciprocity were investigated through the 9-item punishment sub-scale of the Personal Norms of Reciprocity (PNR) measure (Perugini et al., 2003).

### Experimental Design

We examined bargaining behavior using the Ultimatum (Figure 1) (Güth et al., 1982) and Dictator Games (Figure 1) (Kahneman et al., 1986) (∼15 min, counterbalanced across participants). In the Dictator Game (DG), the participant decided how much of an endowed sum ($15-25) to share with their partner. To ensure that participants were deceived into believing that their decisions had a social impact, the participant was told their partner was represented by decisions made by past participants in the study, and that their decisions would be used with future participants. In addition, each decision was made by a different partner, resulting in each trial being a one-shot game. This design is used to minimize the concern for reciprocity, reputation or other motives beyond social preferences for fairness while making each choice (Yamagishi et al., 2012). In the Ultimatum Game (UG), participants acted as the proposer in some trials and the responder in other trials. As the proposer, participants chose a split of their endowment; however, they were aware that their counterpart could reject their offer. As a recipient in the UG, participants were presented offers from partners that they could choose to accept or reject. If they chose to reject the offer, neither they nor the proposer made any money for that trial. Although our hypotheses and analyses were not focused on the recipient decisions, we included this condition to make the task more believable by making participants think that their unfair proposals could be rejected. We characterize strategic behavior as behavior that offers lower amounts in DG and generally higher amounts in UG, as this strategy would maximize earnings and minimize the threat of rejection.

**Figure 1.**
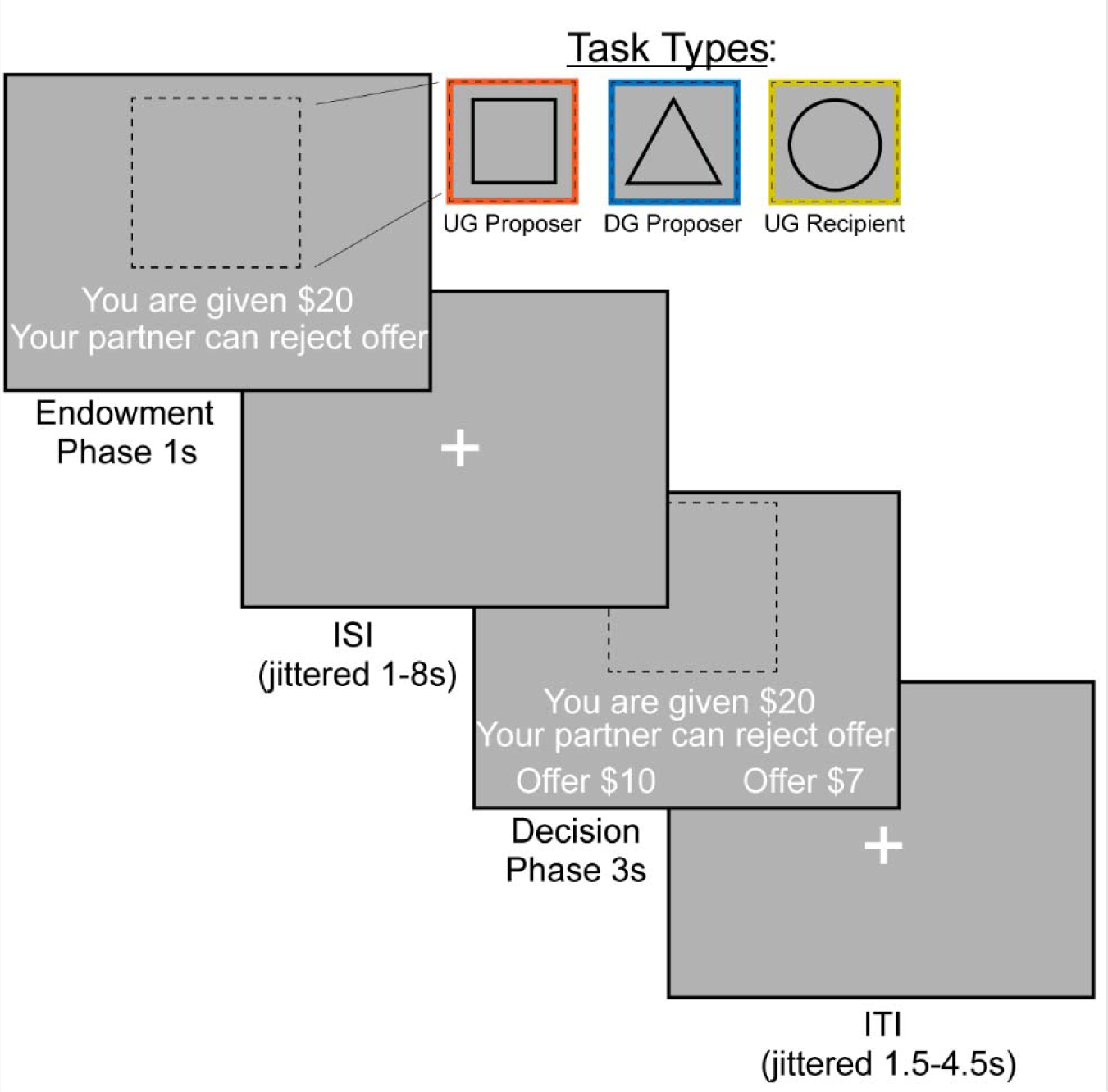
FMRI-based Bargaining tasks to Measure Strategic Behavior Using the Dictator and Ultimatum Games. We operationalized strategic behavior as offering more in the Ultimatum Game and less in the Dictator Game, as this strategy would maximize earnings. During the Endowment phase, the participant learned how much money they were given and which task they would complete. A square indicated that the participant would be acting as the Proposer in the Ultimatum Game or deciding how much money to split with a counterpart. A triangle indicated that the participant would act as the Proposer in the Dictator Game. Finally, a circle indicates that the participant would be the Recipient in the Ultimatum Game, which allowed them to decide whether they would accept or reject an offer given to them. We included the Recipient condition so that participants buy into the manipulation of the threat of punishment during the Ultimatum Game as a proposer. During the Decision Phase, the participant as a proposer decided to offer More or Less to their counterpart. As a recipient, whether to accept or reject the offer.

The experiment consisted of three conditions (Dictator Game-Proposer (DG-P), Ultimatum Game-Proposer (UG-P), Ultimatum Game-Recipient (UG-R)) that were presented in a counterbalanced order. The tasks were administered using PsychoPy (Peirce et al., 2019) across two 7:30 minute runs. Each run consisted of 36 trials, with 12 trials in each condition On each trial, the participant was endowed with a sum of money between $15–$25 and was presented with the type of trial the participant is playing through a cue. If they were acting as the proposer in the DG, they were presented with a triangle. If they were acting as a proposer in the UG, they were presented with a square. Finally, if they were acting as a recipient in the UG they were presented with a circle. Subsequently, the participant experienced an interstimulus interval (ISI) of 1.5-8 seconds, M = 2.7s. During the decision phase as proposer, participants are presented with the option to select a More or Less split. During the decision phase as a recipient, participants have the choice whether to accept or reject the offer. If a participant missed a trial, the screen indicated that they were too slow and recorded a missed trial in the log. Subsequent to each trial, there was a variable duration intertrial interval of 1-4.5 seconds; M = 2.42s.

### Behavioral Data Analysis

Strategic behavior was identified for each participant by calculating how much each person chose to share when there was a threat of punishment versus when there was not a threat of punishment. Specifically, for each participant, we calculated the average proportion of the endowment proposed in UG minus DG. Proportions closer to 0 reflected participants who generally proposed a more even split, whereas proportions closer to 0.5 reflected participants who proposed more unfair offers in DG versus UG. We used this method of measuring strategic behavior rather than pooling hypothetical total earnings (see deviations from pre-registration) as it avoids inferring earnings and simply used the participants’ decisions.

To examine whether participants acted strategically through offering more as a Proposer in the Ultimatum Game condition versus the Dictator Game condition, we used a mixed effect linear model. The regressors included the task (UG-P or DG-P), trial endowment, and the proportion of the endowment the participant offered. While we included the recipient condition (UG-R) so that participants experience offers to understand the threat of punishment as proposers, our main questions do not assess recipient behavior. Nonetheless, as a manipulation check to assess whether participants rejected unfair offers more frequently (i.e., offers with a proportion substantially less than half of the endowment) in the Ultimatum Game as a recipient, we regressed participants’ choices to accept or reject an offer on partner endowment and the proportion offered. Next, we assessed whether there were associations between decisions and measures of social functioning, reward sensitivity, and substance use. Given that both hyper- and hypo-sensitivity to rewards have been linked to substance use (Alloy et al., 2009; Bart et al., 2021; Franken & Muris, 2006), we control for levels of substance use in our data while assessing reward sensitivity. We used correlations between measures (i.e., social functioning, reward sensitivity, and substance use) with the proportions offered in the UG versus DG (i.e., Spitzer et al., 2007). This method of measuring strategic behavior was used rather than pooling hypothetical total earnings (see deviations from pre-registration) as this method avoided inferring earnings and simply used the participants’ decisions. We also conducted exploratory analyses to 1) assess whether there are associations between strategic behavior and reward sensitivity and substance use, and 2) whether there are associations between the individual difference measures and individual conditions (DG-P, UG-P, and UG-R).

We conducted analyses on the included self-report measures to ensure that they were correctly operationalized for further analyses. Since the BAS and SR subscale of the SPSRWD were highly correlated *r*(52) = .71, *p* < .001, we combined them into a single composite measure of reward sensitivity using their combined z-scores. Reward sensitivity scores were binned into deciles to produce an even distribution for subsequent analysis. Finally, because both hyper- and hypo-sensitivity to rewards have been linked to substance use (e.g., Alloy et al., 2009; Bart et al., 2021; Franken & Muris, 2006), we squared the binned composite reward sensitivity scores to create an additional, quadratic measure of aberrant reward sensitivity. In other words, aberrant reward sensitivity explores whether there are consistent patterns across people who are either high or low in reward sensitivity. Next, we found that AUDIT and DUDIT also were correlated *r*(52) = .32, *p* = .02. As a result, we operationalized problematic substance use through z-scoring the responses between the measures and combining them into a single composite z-score of problematic substance use using the same method as described for reward sensitivity. Behavioral data analyses were completed using MATLAB (*MATLAB*, 2022), R (*R Core Team*, 2022), and Python (Van Rossum & Drake, 2009).

### Neuroimaging Data Analyses

Functional images were acquired using a 3T Siemens PRISMA MRI scanner at Temple University. Neuroimaging data were converted to the Brain Imaging Data Structure (BIDS) using HeuDiConv (Halchenko et al., 2024). We applied spatial smoothing with a 5mm full-width at half-maximum (FWHM) Gaussian kernel using FEAT (FMRI Expert Analysis Tool) Version 6.00, part of FSL (FMRIB’s Software Library, www.fmrib.ox.ac.uk/fsl). See the *Supplemental Information* for the full neuroimaging data acquisition and preprocessing pipeline.

Neuroimaging analyses used FSL version 6.0.4 (Jenkinson et al., 2012; S. M. Smith et al., 2004). We used two general linear models with local autocorrelation (Woolrich et al., 2001). Both models included a common set of confound regressors consisting of the six motion parameters (rotations and translations), the first six aCompCor components explaining the most variance, non-steady state volumes, and the framewise displacement (FD) across time. Finally, we used high-pass filtering (128s cut-off) through a set of discrete cosine basis functions.

The first model tested task-based brain activation elicited during the endowment (duration = 1,000 ms) and decision (duration = 3,000 ms) phases and the effect of strategic behavior on brain function during these phases. To do this, we included 6 task-specific regressors (endowment: DG-P, UG-P, and UG-R; decision: DG-P, UG-P, and UG-R), and the same 6 task-specific regressors that we parametrically modulated by the proportion of the offer proposed by the participant. In other words, the parametric modulator measured brain responses to the fairness of the offer proposed. A thirteenth regressor modelled missed trials. By including both parametrically modulated and non-modulated task-based regressors, we were able to investigate the parametric effects while properly controlling for changes in activation across UG and DG conditions.

The second type of model focused on the task-dependent connectivity using the ventral striatum as a seed and areas related to social processing as target regions. To estimate the changes in connectivity resulting from strategic behavior, we used psychophysiological interaction (PPI) analysis (Friston et al., 1997; O’Reilly et al., 2012). Meta-analyses have demonstrated that PPI is able to reliably reveal specific patterns of task-dependent connectivity (D. V. Smith, Gseir, et al., 2016; D. V. Smith, Rigney, et al., 2016; D. V. Smith & Delgado, 2017). Our PPI analysis focused on task-dependent changes in connectivity using the ventral striatum (VS; Oxford-GSK-Imanova atlas) as a seed. Additionally, we used seeds derived from whole-brain analyses (e.g., Inferior Frontal Gyrus and Anterior Insula) to find non-pre-registered target regions in secondary analyses (O’Reilly et al., 2012). The average time course of activation from this seed region was extracted and used as an additional fourteenth regressor. To construct the PPI model, we used the same model described above and added 14 additional regressors (1 regressor for the VS region and 13 regressors for the interaction between the VS region and the task-based regressors), yielding a total of 25 regressors in each seed-based PPI model. Both activation and connectivity models were then run through a fixed effects second level analysis that combined the first and second runs. For participants with missing data, or for runs that were excluded due to head motion, we used a participant’s one good level one run in the group level analyses. For all participants and their combined runs, we used a fixed-effects model.

Group-level analysis focused on activation and connectivity patterns and their associations between bargaining behavior, substance use and BOLD responses, independent of reward sensitivity. The analyses were carried out using FLAME (FMRIB’s Local Analysis of Mixed Effects) Stage 1 (Beckmann et al., 2003; Woolrich et al., 2004). Our group-level model focused on comparisons between the Dictator and Ultimatum Games as a Proposer; these comparisons included covariates to account for reward sensitivity, the second-order polynomial expansion of reward sensitivity (which captures effects tied to aberrant reward sensitivity), substance use, strategic behavior, temporal signal to noise ratio (tSNR) and mean framewise displacement (fd mean). Strategic behavior as a covariate in the group model was identified based on the average proportion offered in UG minus DG for each individual participant. In other words, a participant that was more strategic would have exhibited a larger difference in contributions compared to someone who was less strategic. We also applied two additional models that explored interaction effects. The first interaction model included additional regressors of substance use and reward sensitivity and substance use and aberrant reward sensitivity. The second interaction model included additional regressors of the interaction of strategic behavior and reward sensitivity, and main effects of strategic behavior and aberrant reward sensitivity. We controlled for multiple comparisons through identifying pre-registered regions of interest and by correcting for multiple comparisons across the whole brain using *Z*-statistic images that were thresholded parametrically (Gaussian Random Field Theory) using clusters determined by Z > 3.1 and a (corrected) cluster significance threshold of *P*lJ=lJ0.05 (Flandin & Friston, 2019; Nichols & Hayasaka, 2003; see *Supplemental Information* for more details).

### Deviations from Pre-Registration

Once data collection and analyses began, we made several adjustments based on four issues that were unspecified in our pre-registration. First, we initially specified that we would use the parametric effect of endowment, but not for decisions. For decisions, we expected to use the actual offers selected (High, Low) in our analyses. However, since many participants selected High more often in the UG condition and Low in the DG condition, these regressors had fewer events for comparison. To address this issue, we modeled strategic decisions as parametric effects of offer amount through the difference in the proportions of the endowments offered between DG-P and UG-P. Second, we adjusted the covariates in our group level models due to missing data. Although we originally planned to study Machiavellianism, due to an error in data collection, this survey was not completed by our participants. Next, whereas substance use analyses were not mentioned in the pre-registration, we intended to complete them in accordance with the broader aims and hypotheses of the grant, which are also described in the grant report (Sazhin et al., 2020). Third, we used the (Clithero & Rangel, 2014) (−2, 28, −18) meta-analysis vmPFC coordinates for our mask rather than the mask specified in the pre-registration (Delgado et al., 2016) for greater spatial specificity in our analyses. Fourth, we explored group level models that included the interaction of reward sensitivity, substance use and strategic behavior despite not being initially pre-registered. Taken together, these adjustments from the pre-registration have allowed us to analyze the data more robustly. Our results and discussion take care to differentiate between confirmatory and exploratory results, especially emphasizing differences in our group level models.

## Results

Below, we report results from behavioral analyses, task-based neural activation and connectivity analyses. We begin by presenting results of the behavioral tasks, assessing whether participants made choices as expected, and if their choices relate to self-reported levels of emotional intelligence, attitudes toward rejection, reward sensitivity, and substance use. Next, we examined pre-registered hypotheses examining strategic choices between the dictator and ultimatum games within reward-related and social neural systems (see *Supplemental Information*). Although our pre-registered ROI-based analyses did not support our hypotheses (see *Supplemental Information*), these analyses were followed with a whole-brain analyses that examined activation and connectivity in response to strategic decisions, revealing that elevated IFG and AI activation is associated with strategic decisions. Subsequently, we investigated task-dependent connectivity using the IFG and AI as seeds for potential target regions. These analyses found that IFG-pTPJ connectivity is modulated by elevated strategic decisions. Finally, we present exploratory results that investigate associations between attitudes toward fairness, reward sensitivity, and brain connectivity.

### Strategic Behavior

If participants made higher offers in the Ultimatum Game compared to the Dictator Game, this would indicate that participants were acting most consistently toward maximizing their earnings, thereby exhibiting strategic behavior. Consistent with our expectations, using a mixed effects model for a random intercept, we found that participants (N=54) made more selfish offers in the DG vs. the UG conditions, (B = −0.43, SE = 0.015, *t*(2550) = −28.09, *p* <.001), *see Figure 2),* with the overall model reporting an adjusted *R²* of 0.19. As a manipulation check, we investigated whether participants rejected unfair offers in the recipient condition. A binary logistic regression indicated that participants reject more often with lower offers, (B = 1.72, SE = 0.095, *t*(1252)= 18.06, *p* <.001), with the overall model reporting an adjusted *R²* of 0.50. Next, we explored whether there was a relation between strategic behavior and rejection rate as a function of offer amount as a recipient, finding no significant association, *r*(52) = −.19, *p* =.16. Given that there was no relationship of recipient choices to strategic decisions as proposers, we excluded these measures from subsequent analyses.

**Figure 2.**
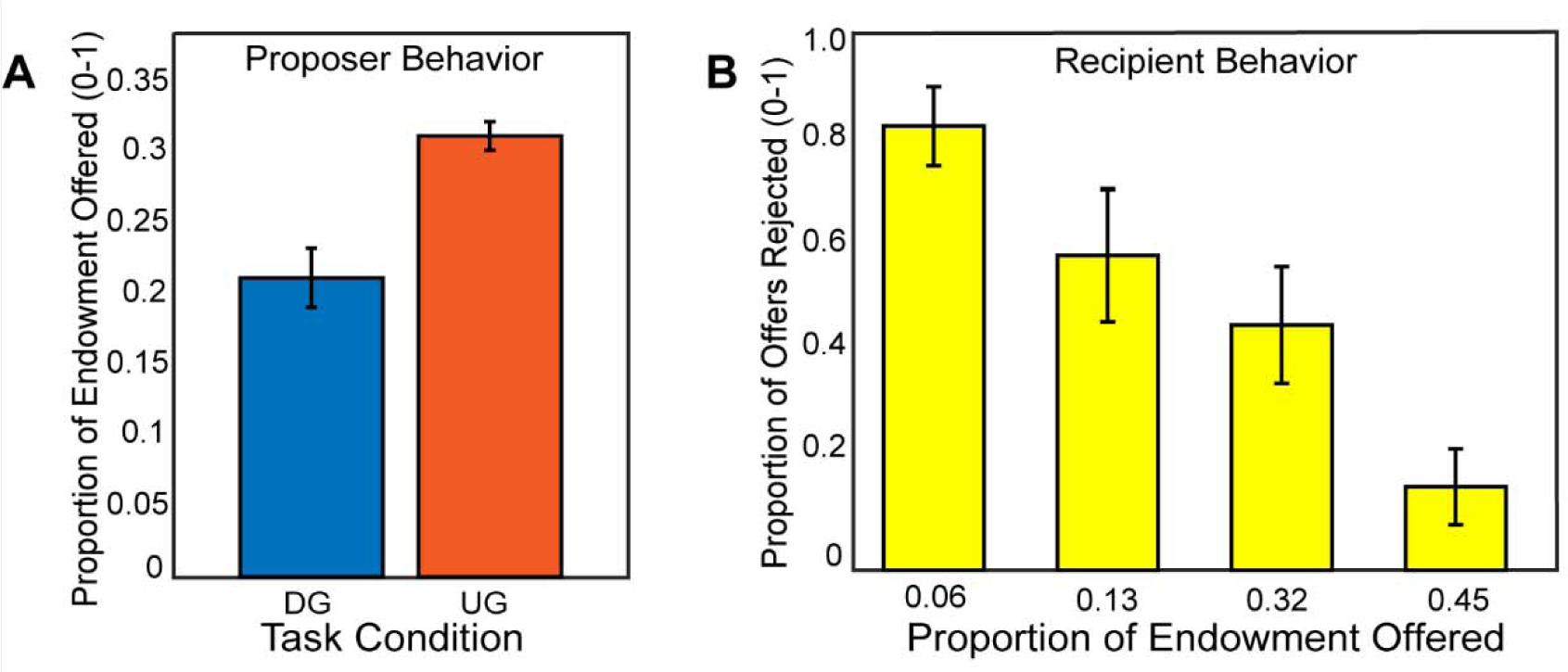
Participants make strategic decisions by offering lower in the Dictator Game versus the Ultimatum Game. In Panel A, we find that participants made higher offers in the Ultimatum Game as a proposer compared to the Dictator Game. In Panel B, we show that participants rejected unfair offers more frequently when they acted as a recipient in the Ultimatum Game. Overall, these behavior results are consistent with our hypotheses and past literature.

**Figure 3.**
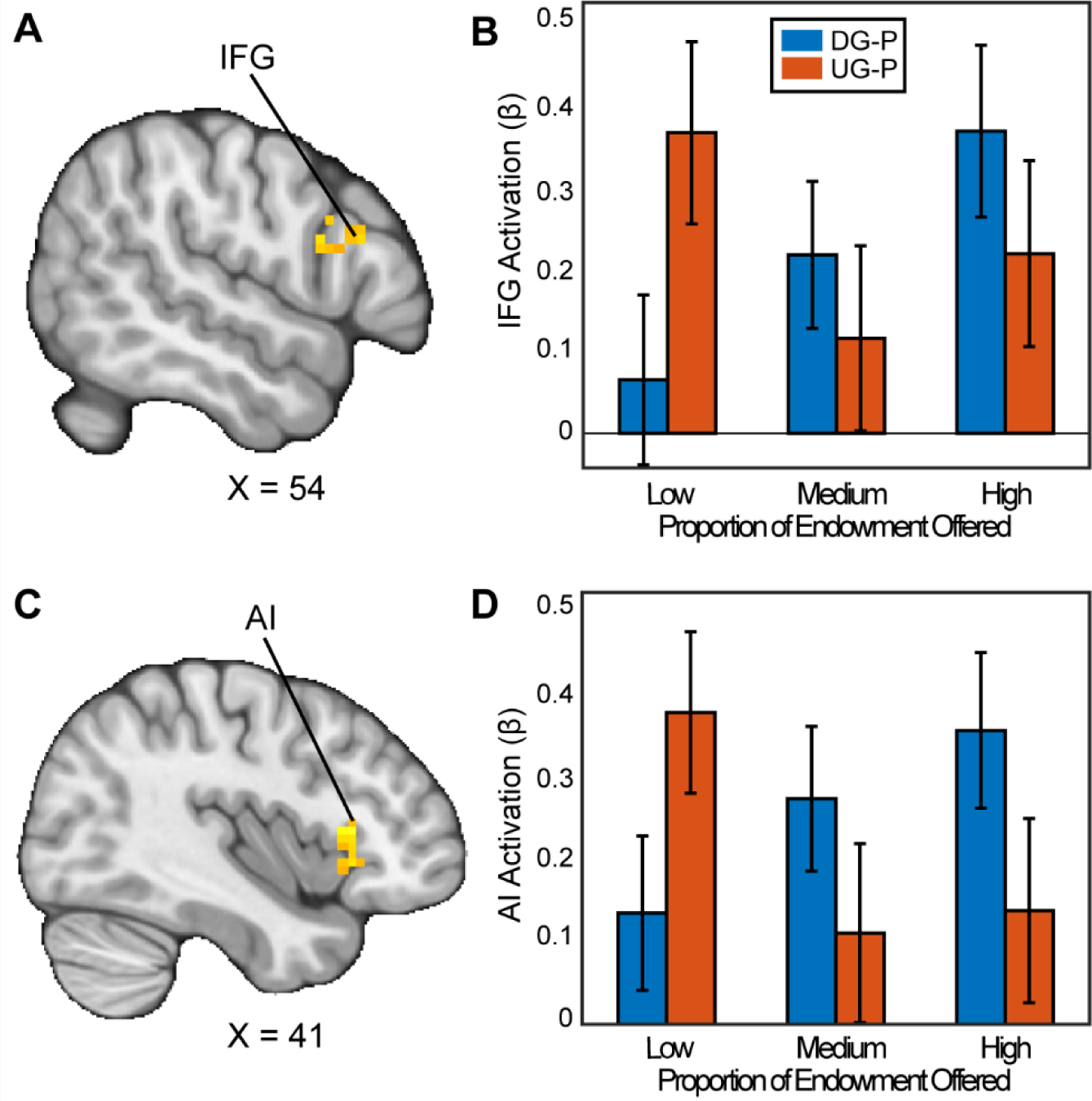
Activation associated with strategic thinking. We conducted a whole-brain analysis to investigate whether there were regions in the brain that differentially responded when acting as a proposer in the DG versus UG conditions. When assessing the parametric effect associated with acting more strategically, Panels A and C reflect regions (Inferior Frontal Gyrus (IFG) (MNIxyz = 52, 16, 22; cluster = 20 voxels, p=0.035, and Anterior Insula (AI) extending into the Orbitofrontal Cortex (OFC) (MNIxyz = 37, 23, 2; cluster = 54 voxels, p<.001 respectively) with significant activation. That is to say, as participants made fairer offers in the DG condition, they experienced stronger activation compared to when they made fairer offers in the UG condition. (Thresholded: https://neurovault.org/images/803473/; Unthresholded: https://neurovault.org/images/803474/). For illustrative purposes, Panels B and D shows the extracted parameter estimates within each region. We note that Z statistic images were thresholded parametrically (Gaussian Random Field Theory) using clusters determined by Z>3.1 and a (corrected) cluster significance threshold of p=.05.

Next, we assessed whether measures of social functioning (N=45) were related to strategic decisions. Several participants had missing questionnaire data, resulting in a smaller dataset for these analyses. Consistent with our hypotheses, individuals scoring higher on the Emotional Intelligence (EI) scale made higher offers as a proposer in the Ultimatum Game, *r*(43) = .35, *p* = .02. Contrary to our hypotheses, we did not find associations between strategic behavior, emotional intelligence, or attitudes toward rejection that met a *p-*value of less than *p*=.05. Inasmuch as there was no effect of strategic behavior and our measures of social functioning as we hypothesized, these measures were excluded from further analyses and used the full dataset of 54 participants for further analyses.

Although we did not expect relations between strategic behavior and measures of reward sensitivity and substance use, we explored whether there were such associations to contextualize any brain relations we may have found with these respective individual difference measures. We did not find any significant associations between reward sensitivity and substance use, and strategic behavior or individual task conditions (DG-P, UG-P, UG-R) that met a threshold of *p* < .05.

### Neural Responses while Making Strategic Decisions

To examine how people make strategic decisions in bargaining situations, we investigated how people propose offers in the Ultimatum Game (UG) versus the Dictator Game (DG). First, we assessed whether there were activation differences between UG and DG conditions, failing to find any significant activation that met *p* = .05 or lower. Next, we assessed if brain responses tracked the fairness of the offers proposed differently between DG and UG. In other words, do participants have differing brain activation when proposing higher proportions of the endowment or lower proportions of the endowment when there is a threat of punishment versus when there is not a threat of punishment?.

Our results indicated that when participants chose to be selfish versus fair in the contrast between the DG and UG as a proposer, there were significant clusters in the Inferior Frontal Gyrus (IFG) (MNIxyz = 51, 24, 24; cluster = 20 voxels, *p*=.035) and a cluster spanning the Anterior Insula (AI), extending into the Orbitofrontal Cortex (OFC) (MNIxyz = 33, 27, −4; cluster = 54 voxels, *p*<.001). We did not find significant activation in the vlPFC or the VS. In the contrast between UG and DG (i.e., choosing to be fair versus unfair), we found a significant cluster in cerebellum (MNIxyz = 30,-82, −36; cluster = 37, *p*<.001). In sum, some of our results successfully replicated past investigations of strategic behavior.

**Table 1:** We incorporated several group level models assessing strategic behavior and reward sensitivity while controlling for substance use. We assessed the interactions of reward sensitivity and strategic behavior and substance use respectively. If there were no interaction effects, we interpreted main effects using the no interaction model. We completed these group level analyses across both activation and PPI models. The PPI model used a pre-registered VS seed, and IFG and AI seeds as derived from our secondary results. The initial group level models were derived from initial hypotheses, though the interaction of reward sensitivity and strategic behavior was an exploratory model driven by our results. Thresholded and unthresholded images are available on Neurovault: https://neurovault.org/collections/15045/

| **<u>Model Type</u>** | **<u>Confirmatory/Exploratory</u>** | **<u>Covariates</u>** |
| --- | --- | --- |
| **No Interactions** | Confirmatory | Strategic Behavior, Substance Use, Reward Sensitivity, Aberrant Reward Sensitivity |
| **Reward Sensitivity x Substance use** | Confirmatory | No Interaction model plus substance use x reward sensitivity, substance use x aberrant reward sensitivity |
| **Reward Sensitivity x Strategic Behavior** | Exploratory | No Interaction model plus strategic behavior x reward sensitivity, strategic behavior x aberrant reward sensitivity |

### Strategic Behavior and Neural Connectivity

Beyond activation patterns, we studied whether task-dependent connectivity patterns related to reward sensitivity and strategic decisions made in the Dictator and Ultimatum games. We included the IFG and AI as seeds because they were derived from the activation of DG versus UG in response to the level of proportion offered. Our group level analyses employed several covariates, including motion-based nuisance regressors, reward sensitivity, substance use, and strategic behavior. We also explored two additional models that investigated the interactions of reward sensitivity, strategic behavior, and substance use respectively.

First, we wanted to examine if strategic behavior as measured by the choices our participants made was associated with brain connectivity. Using the IFG as a seed (MNIxyz = 52, 16, 22), we found that enhanced connectivity with a left rpTPJ target region (Schurz et al., 2017) extending into the SMG (MNIxyz = 50, −68, 35; cluster = 22 voxels, *p* = .008) was modulated by strategic behavior in the Dictator versus Ultimatum Game (see Figure 4). That is to say, selfish participants (i.e.: by making lower proposals in the DG versus UG conditions) experienced enhanced IFG-rpTPJ connectivity contingent on whether or not there was a threat of rejection. Our results suggest that enhanced IFG-rpTPJ connectivity may facilitate the social processing associated with strategic decisions in social contexts. We also examined if connectivity from an AI seed related to strategic situations was modulated by strategic behavior. Using the AI seed (MNIxyz = 33, 27, −4), we found that attenuated connectivity with the neighboring insular cortex (MNIxyz = 50, 6, −1; cluster = 26 voxels, *p* = .003) was modulated by strategic behavior in UG versus DG condition. That is to say, participants who were more selfish when there was no threat of rejection exhibited lower AI-Insula connectivity. Our results suggest that attenuated co-activation of the insular cortex may contribute to making more selfish choices in social contexts.

**Figure 4.**
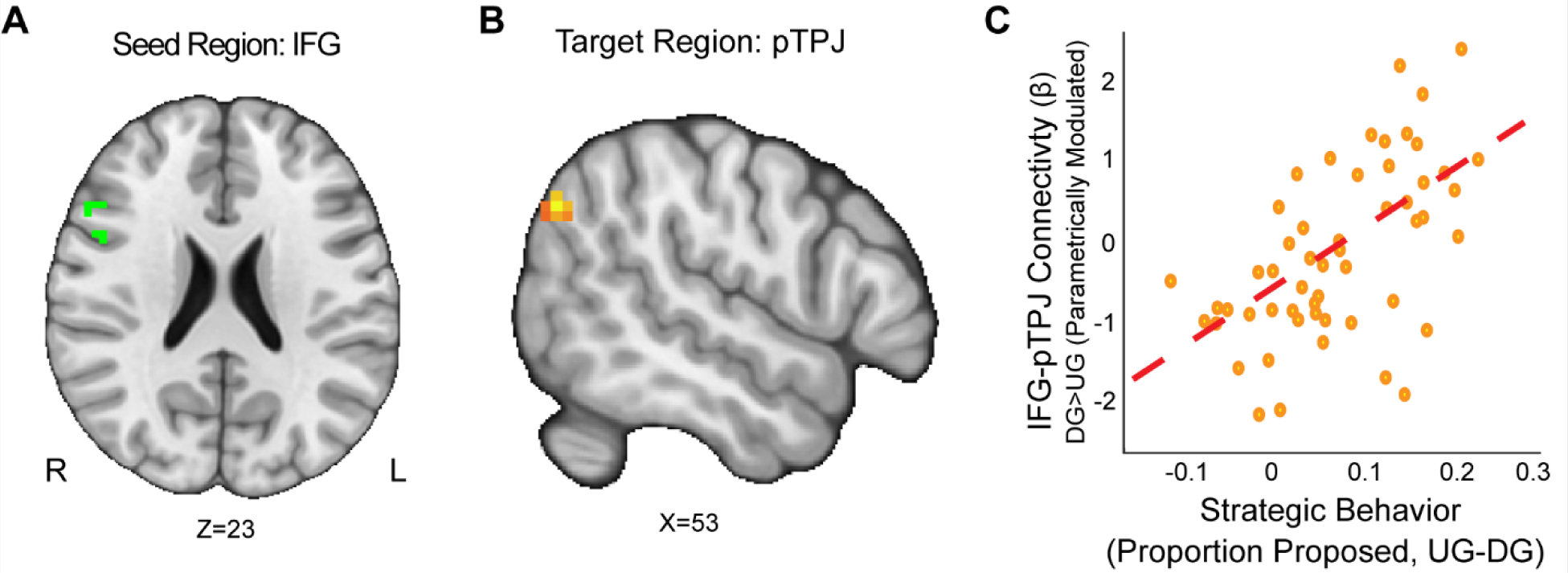
IFG-rpTPJ Connectivity is Modulated by Strategic Behavior. We found that connectivity between an Inferior Frontal Gyrus (IFG) seed (Panel A), and a right pTPJ target (Panel B) was related to elevated strategic behavior (Panel C) (DG > UG) (MNIxyz = 50, −68, 35; cluster = 22 voxels, p = .008).(Thresholded: https://neurovault.org/images/803475/ Unthresholded https://neurovault.org/images/803476/). These results suggest that IFG-right pTPJ connectivity may modulate strategic behavior contingent on whether there is a threat of rejection or not. Participants who experienced elevated IFG-right pTPJ connectivity were also those who were more selfish in DG and offered closer to even splits in UG. For illustrative purposes, we extracted the parameter estimates within each region (Panel C). We note that Z statistic images were thresholded parametrically (Gaussian Random Field Theory) using clusters determined by Z > 3.1 and a (corrected) cluster significance threshold of p=.05 and the images are plotted using radiological view.

### Exploratory Analyses: Anterior Insula-Angular Gyrus Connectivity, Trait Reward Sensitivity, and Strategic Behavior

Next, we explored how the interaction of reward sensitivity and substance use may modulate brain connectivity patterns associated with strategic thinking in bargaining situations.

Investigating how a trait like reward sensitivity may modulate brain responses can reveal an important factor for understanding both behavior and brain relationships. Specifically, we used a model that included interaction covariates of strategic thinking with reward sensitivity and aberrant reward sensitivity. The model also controlled the main effects of strategic behavior, reward sensitivity, aberrant reward sensitivity, and substance use. We included substance use as a controlling variable due to its known relationships with reward sensitivity in psychopathology (Joyner et al., 2019).

We found that the interaction of reward sensitivity and strategic behavior modulated AI-Angular Gyrus connectivity in the UG versus DG condition (Figure 5). That is to say, participants with higher reward sensitivity and attenuated AI-Angular Gyrus connectivity tended to make more strategic choices when there was a threat of rejection relative to when there was not. Moreover, participants with lower reward sensitivity *and* enhanced AI-Angular Gyrus connectivity tended to make more strategic choices when there was a threat of rejection compared to when there was not. These exploratory results suggest that the combination of strategic decisions and a person’s trait reward sensitivity together may modulate connectivity patterns in social situations requiring strategic thinking.

**Figure 5.**
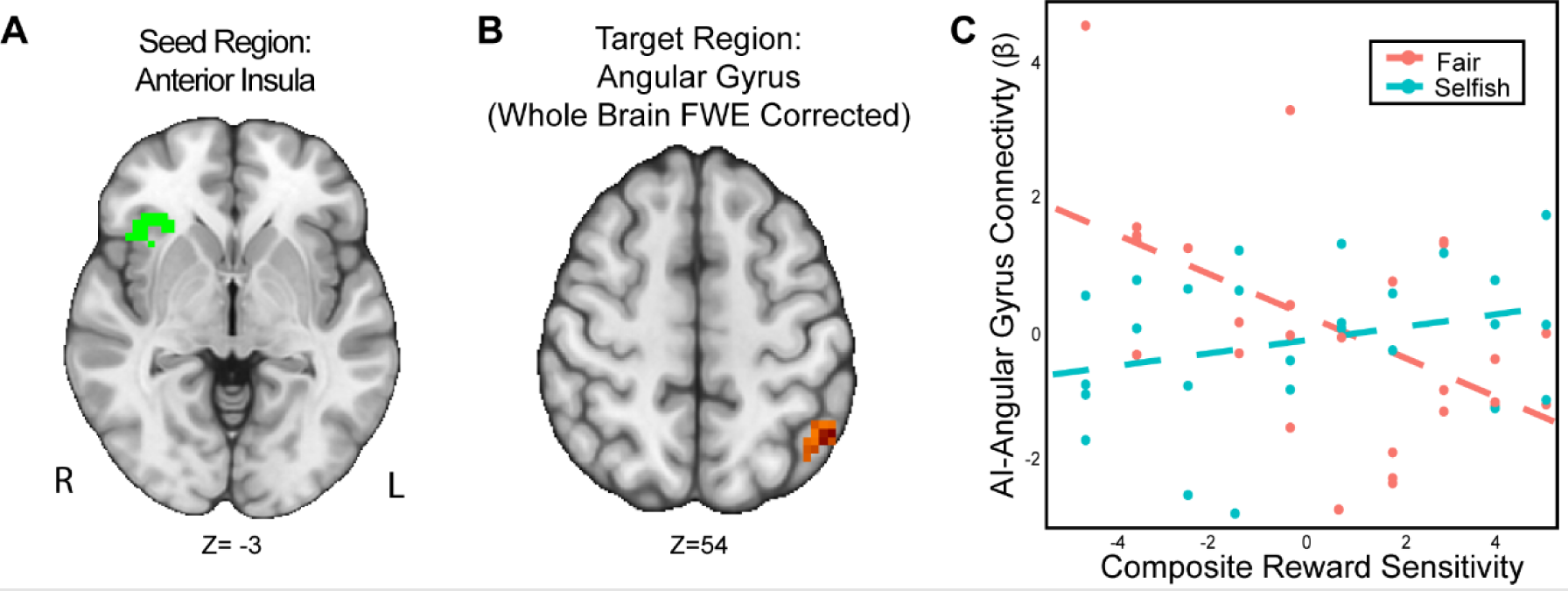
The interaction of reward sensitivity and strategic behavior modulated AI – Angular Gyrus connectivity in social situations requiring strategic thinking. We conducted a whole-brain analysis exploring the interaction of trait reward sensitivity and strategic behavior. We found that higher reward sensitivity is associated with 1) more strategic behavior and 2) elevated task-dependent changes in connectivity between AI (Panel A) and the Angular Gyrus (MNI; xyz = −47,-56, 54; cluster = 23 voxels, p = .005). Conversely, for participants with low reward sensitivity, we found that their AI-Angular connectivity is lower as they exhibit strategic behavior. For illustrative purposes (Panel C), we used a median split to indicate the relation of reward sensitivity and strategic behavior. Next, we extracted the parameter estimates within each region (Panel C). We note that Z statistic images were thresholded parametrically (Gaussian Random Field Theory) using clusters determined by Z > 3.1 and a (corrected) cluster significance threshold of p=.05 and the images are plotted using radiological view. See images here:(Thresholded: https://neurovault.org/images/803477/; Unthresholded: https://neurovault.org/images/803482/).

## Discussion

This study investigated how relations between strategic behavior in bargaining situations and reward responses correspond to patterns of brain activation and connectivity. First, the behavioral results are consistent with past work suggesting that participants act strategically in bargaining situations through acting fairly when there is a threat of rejection (e.g., Ultimatum Game; UG) while keeping more for themselves when there is not a threat of rejection (Dictator Game; DG) (Charness & Gneezy, 2008). Second, the neuroimaging analyses revealed that strategic behavior in the Dictator versus Ultimatum Games evoked activation in the Inferior Frontal Gyrus (IFG) and Anterior Insula (AI), results that were consistent with past investigations (i.e., Spitzer et al., 2007). Our analyses also indicated that elevated IFG-rTPJ connectivity was related to enhanced strategic behavior and attenuated AI-Angular Gyrus connectivity was modulated by the interaction of reward sensitivity and strategic behavior. Taken together, whether people choose to be fair or selfish in bargaining situations is associated with pattern of neural connectivity, which in turn may depend on a person’s trait reward sensitivity.

This work fits in with past literature suggesting that norm compliance is regulated by cortical activation. Although we did not find activation during UG versus DG in the pre-registered regions of interest, whole brain analyses revealed activation in the right IFG and AI as participants made strategic decisions, replicating previous work (Spitzer et al., 2007; Zheng & Zhu, 2013). Next, both IFG and AI activation has been observed in other decision-making contexts. For example, FeldmanHall and colleagues reported AI activation during moral decision making (FeldmanHall et al., 2014). In addition, other work has shown that increased activation in the anterior insula in a trust task is associated with inequity aversion (van Baar et al., 2019; FeldmanHall et al., 2014). Further, our results are consistent with stimulation-based research that found elevated right dlPFC area activation corresponded to more strategic behavior (Knoch et al., 2006; Ruff et al., 2013; Strang et al., 2015) and inhibition of dlPFC activity diminished strategic choices (Müller-Leinß et al., 2018; Zinchenko et al., 2021). In sum, our findings are consistent with the IFG and AI being involved in norm compliance decisions.

Additionally, the results extend on past literature through investigating how reward processes and cortical connectivity interact with strategic behavior. The results indicate that elevated IFG-rpTPJ connectivity is associated with increased strategic behavior, whereas attenuated AI-Angular Gyrus connectivity is modulated by the interaction of reward sensitivity and strategic behavior. Although recent work has shown that the dlPFC and rpTPJ regulate norm compliance in the UG and DG (Gianotti et al., 2018), and that the right TPJ does not necessarily yield greater generosity (Brethel-Haurwitz et al., 2022), the results indicate that strategic decision making in social situations modulates the connectivity between the dlPFC and TPJ. Understanding how connectivity modulates strategic decisions is a critical component of characterizing how the TPJ and dlPFC may be regulated during decision making, with the TPJ potentially orienting the IFG toward changes to social context and thus greater opportunities to be strategic. Additionally, when including reward sensitivity as a covariate, the results indicated that people with varying levels of trait reward sensitivity respond to strategic decisions differently. Specifically, people with low reward sensitivity are more strategic with decreasing AI-Angular Gyrus connectivity, whereas people with higher reward sensitivity are more strategic with increasing AI-Angular Gyrus connectivity.

It has been previously found that reward sensitivity is associated with risky behavior (Scott-Parker & Weston, 2017), higher Machiavellianism (Birkás et al., 2015), and more strategic behavior (Scheres & Sanfey, 2006). This yields an interpretation that reward sensitivity could be a factor in guiding norm compliance in social situations as people with higher reward sensitivity may be more motivated to maximize their own rewards. Specifically, reward sensitivity may modulate strategic decisions by increasing the degree people are self-oriented, and their willingness to take risk even at the potential of being rejected in a bargaining situation. Thus, AI-Angular Gyrus connectivity may modulate how people experience opportunities to cooperate and defect, which could affect how people employ social heuristics in bargaining situations We speculate that among self-interested people who aim to maximize earnings, reward sensitivity may modulate strategic decisions through increasing attentional processes to changes in social context through AI-Angular Gyrus connectivity. Specifically, connectivity between the AI-Angular Gyrus may serve as a mechanism for overriding fairness norms to share evenly with their partner by orienting people to changes in social context. This process could be driven through bottom-up attention, or through top-down cognitive mechanisms.

Specifically, the angular gyrus is implicated in bottom-up attentional processing (Cabeza et al., 2012; Seghier, 2013), interpreting contextual information (Carter & Huettel, 2013; Ramanan et al., 2018), and social cognition (Numssen et al., 2021). The AI integrates fairness, empathy, and cooperation (Cheng et al., 2017; Lamm & Singer, 2010). Given this, it is plausible that AI-Angular Gyrus connectivity could help bottom-up orientation of changes in context affecting social norms. Alternatively, AI engagement could reflect differences in top-down cognitive control among participants (Sridharan et al., 2008), and AI-angular gyrus connectivity may reflect top-down orientation to the changes in social context. Additionally, AI-Angular Gyrus connectivity may be modulated by reward sensitivity. Reward sensitivity is associated with risky behavior (Scott-Parker & Weston, 2017), higher Machiavellianism (Birkás et al., 2015), and more strategic behavior (Scheres & Sanfey, 2006). Thus, AI-Angular Gyrus connectivity may modulate how people experience opportunities to cooperate and defect, which could affect how people employ social heuristics in bargaining situations.

One interpretation is that people with higher reward sensitivity may be more motivated in the task and may be more likely to defect in bargaining tasks. Increased deliberation may, in turn, override default fairness norms. This deliberative process may modulate bottom-up attention or contextual orienting in the Angular Gyrus, or top-down cognitive processing in the AI. Our results suggest a nuanced view of AI-Angular Gyrus and IFG-TPJ coupling (Lockwood et al., 2020), indicating that these brain regions do not necessarily reflect altruistic choice on their own (Hutcherson et al., 2015), but may modulate cognitive reward processes while making social decisions. Additionally, we speculate that our results reflect that downregulation of bilateral TPJ activation and AI deactivation (FeldmanHall et al., 2014) interacts with trait reward sensitivity.

Specifically, our findings may provide insight into how people with aberrant levels of reward sensitivity respond to strategic decisions in bargaining situations. The results indicated that people with lower reward sensitivity had higher AI-Angular Gyrus connectivity when being less strategic, whereas people with higher reward sensitivity had higher connectivity when being more strategic. If higher AI-Angular Gyrus connectivity is a reflection of increased motivation among participants, the results suggest that trait reward sensitivity may inform strategic behavior and how people employ social heuristics to be fair or selfish. Specifically, people who have higher reward sensitivity may need to have greater AI-Angular gyrus connectivity to be more strategic compared to people who have lower reward sensitivity. Additionally, since aberrant reward sensitivity is a predictor for elevated substance use, investigating how reward sensitivity modulates brain processes in social contexts could provide insight into how people make decisions resulting in substance use (Bart et al., 2021; Heilig et al., 2016; Wyngaarden et al., 2023).”

Although our work has found that strategic behavior is modulated by both AI-Angular Gyrus and IFG connectivity with the TPJ, and reward sensitivity, we acknowledge that our study has several limitations that merit discussion. First, although the results suggest bilateral TPJ connectivity and strategic behavior, we do not infer specificity in lateralization. Past investigations suggest mixed findings (Carter et al., 2012; Coricelli & Nagel, 2009; Saxe & Kanwisher, 2003) as to the roles of the right and left TPJ, and we judged that exploring these results further was beyond the scope of this paper. Additionally, we acknowledge that connectivity analyses are not causal or directional with respect to inference despite identifying the IFG and AI as seeds and the temporoparietal junction as target. Further, since strategic behavior as a proposer was not related to recipient choices, we judged that these results are beyond the scope of this investigation. A possible future direction includes evaluating AI-Angular Gyrus and IFG-TPJ connectivity patterns, associations with reward sensitivity, and their relations with recipient decisions in the Ultimatum Game.

Second, we acknowledge that fMRI analysis techniques carry elevated risk of Type II errors. The results reported in the manuscript are a product of whole-brain analyses which were conservatively thresholded to control for multiple comparisons whereas our confirmatory ROI-based analyses registered null results. In line with recommended practices (Gentili et al., 2021), we pre-registered and conducted ROI-based analyses to increase power to detect activation and connectivity by limiting multiple comparisons (Poldrack et al., 2007). Secondary whole-brain analyses naturally follow ROI analyses if appropriately thresholded (Poldrack, 2007; Szycik et al., 2009) and were reported accordingly. Nonetheless, conducting brain-wide association tests with individual difference measures may be underpowered (Marek et al., 2022). Thus, while the sample included people with high and low reward sensitivity and conducted rigorous test-retest with SR and BIS/BAS to ensure that participants were consistent across these measures, we acknowledge that relations with reward sensitivity should be considered exploratory in nature. Future analyses could examine how reward sensitivity modulates brain responses using multivariate methods to improve effect size estimation (Reddan et al., 2017) with canonical correlation analysis (Zhuang et al., 2020), multivariate pattern analysis (Kragel & LaBar, 2015) or machine learning algorithms to assess neural signatures (Wager et al., 2013) of bargaining.

Third, we note that relations with social context, reward sensitivity, and brain connectivity could be studied more extensively in a clinical population to assess how these relations are modulated by substance use and manic-depressive symptoms. Whereas we were able to control for levels of substance use to account for reward sensitivity effects (Joyner et al., 2019), the sample had too limited variability in substance use to make inferences about how substance use may contribute to maladaptive strategic decisions. Additionally, while we assessed strategic behavior, we did not assess it in a dynamic context. As social contexts increase exploration and obtained rewards (Plate et al., 2023), a fruitful future direction could investigate how brain responses to changes over time reflect social decisions.

A final notable limitation was that we did not find evidence that suggests ventral striatal activation or connectivity is related to strategic behavior. Past investigations suggested that higher VS activation was associated with more strategic behavior (Spitzer et al., 2007), with more unfair offers in UG being associated with higher VS activation (Chen et al., 2017). Thus, it is possible that the lack of VS activation was due to participants not finding the differences in offers sufficiently salient, or not being sufficiently incentivized by the small differences in rewards between UG and DG, or potentially that we were underpowered to detect these effects. Alternatively, some individuals may have increased VS activation that may be responding to prosociality, when giving more money to their partner. Across the DG, studies have found increased VS activation for keeping more for oneself (Tricomi et al., 2010), and for sharing with others (Moll et al., 2006). In the UG, the VS tracked inequity in both prosocial and individualistic people (Haruno et al., 2014). Thus, it is possible that in our sample we had individuals that had higher VS activation and acted least strategically toward maximizing their own earnings. Future studies may be able to investigate if there is higher VS activation between people who maximize earnings for themselves or for others across the UG and DG tasks.

Despite the limitations, our findings indicate that strategic decisions in social contexts are associated with elevated IFG-TPJ connectivity and that AI-Angular Gyrus connectivity while making strategic decisions is modulated by trait reward sensitivity. These results provide greater insights into how reward processes interact with social decisions, involving brain processes that appraise the roles of other people while making choices. Since aberrant reward sensitivity is a major mechanism in substance use and depressive and bipolar disorders, investigating how reward sensitivity modulates brain processes during social contexts could provide considerably more understanding into how people make maladaptive decisions resulting in substance use (Bart et al., 2021; Heilig et al., 2016; Wyngaarden et al., 2024). Such work could help identify people at risk for substance use disorders and help develop interventions for people with aberrant reward patterns.

## Supporting information

Supplemental Methods and Results

## Acknowledgments

This work was supported in part by grants from the National Institute of Mental Health (R01-MH123473 and R01-MH126911 to LBA), the Eunice Kennedy Shriver National Institute of Child Health and Human Development (R21-HD093912 to JMJ), the National Institute on Aging (RF1-AG067011 to DVS), and the National Institute on Drug Abuse (R03-DA046733 to DVS), and also a fellowship from the Temple Public Policy Lab (to JMJ).

## Data and code availability

Analysis code related to this project can be found on GitHub: (https://github.com/DVS-Lab/istart-ugdg). Thresholded and unthresholded statistical maps are located on https://neurovault.org/collections/15045/. In addition, all raw data is made available on OpenNeuro (https://openneuro.org/datasets/ds004920/versions/1.1.1).

