## Supplemental Methods and Results for "Trait Reward Sensitivity Modulates Connectivity with the Temporoparietal Junction and Anterior Insula during Strategic Decision Making"

### **Neuroimaging Data Acquisition**

Functional images were acquired using a 3T Siemens PRISMA MRI scanner. Bold Oxygenation Level-Dependent (BOLD) sensitive functional images were acquired using a gradient echo-planar imaging (EPI) sequence (240 mm in FOV, TR = 1,750 ms, TE = 29 ms, voxel size of 3.0 x 3.0 x 3.0 mm3, flip angle = 74°, interleaved slice acquisition). Each of the two runs included 225 functional volumes. We also collected single-band reference images with each functional run of multi-band data to improve motion correction and registration. To facilitate anatomical localization and co-registration of functional data, a high-resolution structural scan was acquired (sagittal plane) with a T1-weighted magnetization=prepared rapid acquisition gradient echo (MPRAGE) sequence (224 mm in FOV, TR = 2,400 ms, TE = 2.17 ms, voxel size of 1.0 x 1.0 x 1.0 mm3, flip angle 8°). In addition, we also collected a B0 fieldmap to unwarp and undistort functional images (TR: 645 ms; TE1: 4.92 ms; TE2: 7.38 ms; matrix 74×74; voxel size: 2.97×2.97×2.80 mm; 58 slices, with 15% gap; flip angle: 60°).

#### **Preprocessing of Neuroimaging Data**

Neuroimaging data were converted to the Brain Imaging Data Structure (BIDS) using HeuDiConv version 0.9.0 (Halchenko et al., 2019). Results included in this manuscript come from preprocessing performed using fMRIPrep 20.2.3 (Esteban et al., 2018, 2019), which is based on Nipype 1.4.2 (K. Gorgolewski et al., 2011; K. J. Gorgolewski et al., 2017). The details described below are adapted from the fMRIPrep preprocessing details; extraneous details were omitted for clarity. Head motion was calculated to determine exclusions. Excess head motion was defined where at least one run was a motion outlier, characterized by either fd mean exceeding 1.5 times the interquartile range above the 75th or tsnr values lower than 1.5 times the lower bound minus the 25th percentile per neuroimaging data quality measures from MRIQC).

**Anatomical data preprocessing.** The T1-weighted (T1w) image was corrected for intensity non-uniformity (INU) with ‘N4BiasFieldCorrection’, distributed with ANTs 2.3.3, and used as T1w-reference throughout the workflow. The T1w-reference was then skull-stripped with a Nipype implementation of the `antsBrainExtraction.sh` workflow (from ANTs), using OASIS30ANTs as target template. Brain tissue segmentation of cerebrospinal fluid (CSF), white-matter (WM), and gray-matter (GM) was performed on the brain-extracted T1w using `fast` (FSL 5.0.9). Volume-based spatial normalization to one standard space (MNI152NLin2009cAsym) was performed through nonlinear registration with `antsRegistration` (ANTs 2.3.3), using brain-extracted versions of both T1w reference and the T1w template. The following template was selected for spatial normalization: ICBM 152 Nonlinear Asymmetrical template version 2009c (TemplateFlow ID: MNI152NLin2009cAsym).

**Functional data preprocessing.** For each BOLD run, the following preprocessing steps were performed. First, a reference volume and its skull-stripped version were generated by aligning and averaging 1 single-band references (SBRefs). A B0-nonuniformity map (or fieldmap) was estimated based on a phase-difference map calculated with a dual-echo GRE (gradient-recall echo) sequence, processed with a custom workflow of SDCFlows inspired by the `epidewarp.fsl` script (http://www.nmr.mgh.harvard.edu/~greve/fbirn/b0/epidewarp.fsl) and further improvements in HCP Pipelines. The fieldmap was then co-registered to the target EPI (echo-planar imaging) reference run and converted to a displacements field map (amenable to registration tools such as ANTs) with FSL's `fugue` and other SDCflows tools. Based on the estimated susceptibility distortion, a corrected EPI (echo-planar imaging) reference was calculated for a more accurate co-registration with the anatomical reference. The BOLD reference was then co-registered to the T1w reference using `flirt` (FSL 5.0.9) with the boundary-based registration cost-function. Co-registration was configured with nine degrees of freedom to account for distortions remaining in the BOLD reference. Head-motion parameters with respect to the BOLD reference (transformation matrices, and six corresponding rotation and translation parameters) are estimated before any spatiotemporal filtering using `mcflirt`.

BOLD runs were slice-time corrected using `3dTshift` from AFNI 20160207. First, a reference volume and its skull-stripped version were generated using a custom methodology of fMRIPrep. The BOLD time-series (including slice-timing correction when applied) were resampled onto their original, native space by applying a single, composite transform to correct for head-motion and susceptibility distortions. These resampled BOLD time-series will be referred to as preprocessed BOLD in original space, or just preprocessed BOLD.

The BOLD time-series were resampled into standard space, generating a preprocessed BOLD run in MNI152NLin2009cAsym space. First, a reference volume and its skull-stripped version were generated using a custom methodology of fMRIPrep. Several confounding time-series were calculated based on the preprocessed BOLD: framewise displacement (FD), and three region-wise global signals. FD was computed using two formulations following Power (absolute sum of relative motions) and Jenkinson (relative root mean square displacement between affines). FD is calculated for each functional run using its implementations in Nipype. The three global signals are extracted within the CSF, the WM, and the whole-brain masks.

Additionally, a set of physiological regressors were extracted to allow for component-based noise correction (CompCor). Principal components are estimated after high-pass filtering the preprocessed BOLD time-series (using a discrete cosine filter with 128s cut-off) for anatomical component correction (aCompCor). For aCompCor, three probabilistic masks (CSF, WM and combined CSF+WM) are generated in anatomical space. The implementation differs from that of Behzadi et al. in that instead of eroding the masks by 2 pixels on BOLD space, the aCompCor masks are subtracted from a mask of pixels that likely contain a volume fraction of GM. This mask is obtained by thresholding the corresponding partial volume map at 0.05, and it ensures components are not extracted from voxels containing a minimal fraction of GM. Finally, these masks are resampled into BOLD space and binarized by thresholding at 0.99 (as in the original implementation). Components are also calculated separately within the WM and CSF masks. For each CompCor decomposition, the k components with the largest singular values are retained, such that the retained components' time series are sufficient to explain 50 percent of variance across the nuisance mask (CSF, WM, combined, or temporal). The remaining components are dropped from consideration. The head-motion estimates calculated in the correction step were also placed within the corresponding confounds file. All resamplings can be performed with a single interpolation step by composing all the pertinent transformations (i.e., head-motion transform matrices, susceptibility distortion correction when available, and co-registrations to anatomical and output spaces). Gridded (volumetric) resamplings were performed using `antsApplyTransforms` (ANTs), configured with Lanczos interpolation to minimize the smoothing effects of other kernels. Many internal operations of fMRIPrep use Nilearn 0.6.2, mostly within the functional processing workflow. For more details of the pipeline, see the section corresponding to workflows in fMRIPrep's documentation (https://fmriprep.readthedocs.io/en/latest/workflows.html).

Next, we applied spatial smoothing with a 5mm full-width at half-maximum (FWHM) Gaussian kernel using FEAT (FMRI Expert Analysis Tool) Version 6.00, part of FSL (FMRIB’s Software Library, www.fmrib.ox.ac.uk/fsl). Non-brain removal using BET (Smith et al., 2002) and grand mean intensity normalization of the entire 4D dataset by a single multiplicative factor were also applied.

### **Adjusting For Multiple Comparisons**

We used both region of interest (ROI) and whole-brain analyses to correct for multiple comparisons and help balance Type I and Type II errors. When pre-registered and conducted properly (Gentili et al., 2021; Kriegeskorte et al., 2009), ROI-based analyses increase power to detect activation and connectivity by limiting multiple comparisons (Poldrack et al., 2007). Our primary analyses therefore used pre-registered ROI-based analyses. We used the Harvard-Oxford Atlas to make masks for the dlPFC, mPFC (i.e., paracingulate gyrus), SPL, ACC, and Insula. We used the Oxford-GSK-Imanova and Striatal Connectivity Atlases to define the nucleus accumbens mask. Next, we used the Mars TPJ Atlas for our right pTPJ mask. Finally, for the vmPFC mask, we used the Clithero and Rangel, 2013 meta-analysis coordinates (-2, 28, -18) and a 5mm sphere.

Although ROI-based analyses can increase sensitivity, they necessarily ignore effects outside of hypothesized regions. To address this important limitation, we followed each ROI-based analysis with a secondary analysis that corrected for multiple comparisons across the whole brain (Poldrack, 2007; Szycik et al., 2009). Specifically, all z-statistic images were thresholded and corrected for multiple comparisons using an initial cluster-forming threshold of z > 3.1 followed by a whole-brain corrected cluster-extent threshold of p < 0.05, as determined by Gaussian Random Field Theory (Cao & Worsley, 2001; Eklund et al., 2016; Flandin & Friston, 2019; Nichols & Hayasaka, 2003). Additionally, we investigated exploratory, non-pre-registered group level models using whole-brain analyses. Both secondary, and exploratory analyses reported coordinates using the center of gravity (CoG). In line with recommended practices (Gentili et al., 2021), we pre-registered our initial ROIs, performed whole-brain analyses, and provided voxel-wise unthresholded maps of effect sizes on Neurovault.

### **Supplemental Results**

In our pre-registration, we specified hypotheses related to strategic decision making and resultant activation and task-dependent connectivity in several regions-of-interest during the endowment and decision phases. We did not find evidence supporting activation or connectivity in these regions-of-interest. We describe below the regions we investigated during the endowment and decision phases of the Ultimatum Game (UG) and Dictator Game (DG).

#### **Neural Responses during Endowment**

Our first goal for our neuroimaging analyses was to examine the response to endowment. We hypothesized that responses in the ventral striatum would increase as a function of the endowment size. We conducted an ROI-based confirmatory analysis, expecting activation in the VS and the vmPFC. We expected that VS and vmPFC responses would vary based on the expectation of how much of the endowment would subsequently be kept in the decision phase, with the greatest amounts in DG-P, moderate in UG-P, and lowest in UG-R. Contrary with our hypotheses, we did not find a significant difference in vmPFC activation during the endowment phase between the DG-P, UG-P, and UG-R conditions, using a one-way ANOVA (2,159) *F* = 2.40 *p* = 0.09. Next, we did not find significant activation in the VS during the endowment phase when we used a one-way ANOVA to compare activation between the DG-P, UG-P, and UG-R task conditions *F*(2,159) = 1.10 *p* = 0.34 as participants received greater proportions of the endowment.

Next, we investigated whether there were associations between reward sensitivity, neural activation, and the levels of endowment presented. We expected that the response in the VS and vmPFC would positively vary based on the endowment. Such an association would reveal if there were moderating variables in VS or vmPFC activation as participants are endowed with higher levels of money when there is a threat of rejection (UG) versus when there is not (DG). Neither confirmatory ROI-based analyses nor secondary whole-brain analyses found an association between the level of activation in the vmPFC or VS, amount of endowment, and substance use or reward sensitivity.

### **Neural Responses to Strategic Decisions**

We initially conducted confirmatory analyses to investigate how the brain responds as people exhibit strategic behavior through offering fairer offers in the Ultimatum Game (UG) versus the Dictator Game (DG). We hypothesized that we would find activation in areas previously examined (Spitzer et al., 2007), and brain areas associated with social processing, specifically the Anterior Cingulate Cortex (ACC), Superior Parietal Lobule (SPL), and posterior temporoparietal junction (rpTPJ). We conducted a ROI analysis in each of these regions and we did not find any significant activation that met *p* = .05 or lower.

### **Ventral Striatal Connectivity in Response to Strategic Decisions**

First, we conducted confirmatory analyses to examine our pre-registered ROI-based hypotheses using the ventral striatum as a seed. During the decision phase for the proposer conditions, we expected to find that ventral striatal responses to situations requiring strategic thinking (UG-P > DG-P) would be associated with enhanced connectivity with regions modulated by social information. To test this hypothesis, we conducted a PPI analysis using the VS as a seed region, and we focused on connectivity with a priori target regions (vmPFC, mPFC as defined by paracingulate gyrus, rpTPJ, and the SPL). Contrary to our hypotheses, we did not find any significant connectivity between a VS seed and our expected regions of interest that survived multiple comparisons that met *p* = .05 or lower.
